# Syndecan-4 Proteoliposomes Enhance Revascularization in a Rabbit Hind Limb Ischemia Model of Peripheral Ischemia

**DOI:** 10.1101/2022.09.21.508898

**Authors:** Andrew D Sligar, Gretchen Howe, Julia Goldman, Patricia Felli, Almudena Gómez-Hernández, Eri Takematsu, Austin Veith, Shubh Desai, William J. Riley, Rohan Singeetham, Lei Mei, Gregory Callahan, David Ashirov, Richard Smalling, Aaron B. Baker

**Author notes:** Correspondence to: Aaron B. Baker, Ph.D., University of Texas at Austin, Department of Biomedical Engineering, 1 University Station, BME 5.202D, C0800, Austin, TX 78712.

## Abstract

Regenerative therapeutics for treating peripheral arterial disease are an appealing strategy for creating more durable solutions for limb ischemia. In this work, we performed preclinical testing of an injectable formulation of syndecan-4 proteoliposomes combined with growth factors as treatment for peripheral ischemia delivered in an alginate hydrogel. We tested this therapy in an advanced model of hindlimb ischemia in rabbits with diabetes and hyperlipidemia. Our studies demonstrate enhancement in vascularity and new blood vessel growth with treatment with syndecan-4 proteoliposomes in combination with FGF-2 or FGF-2/PDGF-BB. The effects of the treatments were particularly effective in enhancing vascularity in the lower limb with a 2-4 increase in blood vessels in the treatment group in comparison to the control group. In addition, we demonstrate that the syndecan-4 proteoliposomes have stability for at least 28 days when stored at 4°C to allow transport and use in the hospital environment. In addition, we performed toxicity studies in the mice and found no toxic effects even when injected at high concentration. Overall, our studies support that syndecan-4 proteoliposomes markedly enhance the therapeutic potential of growth factors in the context of disease and may be promising therapeutics for inducing vascular regeneration in peripheral ischemia.

## Introduction

Peripheral arterial disease (PAD) is the narrowing of arteries in the extremities, most commonly in the lower limbs. This disorder is a rapidly expanding medical issue worldwide due to increased incidence of metabolic disorders, obesity, and diabetes.^1^ In the U.S., it is estimated that approximately 20% of people over 65 years have PAD and that there 8.5 to 12 million patients with PAD.^2^ Worldwide there are greater than 230 million patients with PAD.^3^ Major consequence of PAD is the development of ischemia in the limbs. As such, PAD often manifests as intermittent claudication and, in some cases, as critical limb ischemia (CLI). Ultimately, long term limb ischemia has the potential to lead to nonhealing ulcers, gangrene, enhanced risk for amputation and death.^4^ The fundamental drivers of PAD include the formation of atherosclerotic plaque and microvascular disease. The development of limb ischemia implies that there is an insufficient regenetive response from the patient in terms of developing new blood vessels through the processes of angiogenesis and arteriogenesis. Current treatments for PAD include percutaneous interventions, drug treatments including vasodilators and anti-thrombotics, and surgical bypass. While these methods provide relief for some patients, they do not address the lack of microvascular perfusion of the tissue and often poorly address the diffuse nature of PAD within the lower limb.^5^ In theory, the ideal approach to treating limb ischemia would be the stimulation of angiogenesis and arteriogenesis to enable the restoration of blood flow in the ischemia tissue.^6^ Growth factors that induce blood vessel growth would seem to be an appealing therapeutic option for enhancing vascular regeneration. Among these, FGF-2 and VEGF have been explored in clinical trials for intermittent claudication and critical limb ischemia. However, the outcomes of the clinical trials testing regenerative approaches to treating PAD have been disappointing, showing a lack of efficacy or minimal efficacy for inducing regeneration in human patients.^7,8^ There are likely multiple reasons for these clinical failures, including issues relating to the delivery of the compounds and the non-responsiveness of human patients with long term disease. We and others have shown that there is significant therapeutic resistance to inducing angiogenesis/arteriogenesis due to common co-morbidities that are found with PAD, including diabetes, hyperlipidemia and obesity.^9-14^ In addition, as preclinical studies for PAD are often performed using healthy animals with surgical ligation, these animal models are often poorly predictive of therapeutic efficacy in human patients that have developed peripheral ischemia.

In past studies, we have shown that that diabetes and hyperlipidemia lead cell surface proteoglycans such as syndecan-4 and glypican-1 in animal models and in human patients.^12,13^ In addition, other studies have shown loss of heparan sulfate proteoglycans in diabetes and an increase in heparanase, the human enzyme that degrades heparan sulfate, in diabetes and severe atherosclerotic plaques.^11,13,15,16^ These complex molecules play important role in many aspects of vascular regeneration including stabilizing growth factor/growth factor receptor interactions, regulating solution, protecting growth factors from degradation, and controlling subcellular trafficking of growth factors.^17,18^ Our previous work has shown that syndecan-4 proteoliposomes (S4PLs) dramatically enhance the recovery of perfusion in hind limb ischemia models in both rats and diabetic, hyperlipidemic mice.^10,11,19^ In addition, we have shown this therapy can increase dermal wound healing and angiogenesis in diabetic, hyperlipidemic mice when combined with FGF-2 or PDGF-BB.^9,12^ Mechanistically, we have shown that syndecan-4 proteoliposomes enhance growth factor signaling, increases trafficking of FGF-2 to the nucleus, facilitates receptor/growth factor recycling and modulates macrophages to an M2 phenotype.^10,11,19^ In the study, we have performed translational studies to evaluate the safety and efficacy of S4PLs delivered in an injectable alginate gel in conjunction with growth factors that induce angiogenesis and arteriogenesis in a large animal model of peripheral ischemia. We evaluated the therapy in a rabbit model of limb ischemia recently developed by our group that includes diabetes/hyperlipidemia and exhibits therapeutic resistance to revascularization.^14^ Our studies demonstrate that growth factors with S4PLs markedly improve revascularization in ischemia in this model whereas growth factor therapy alone was ineffective.

## Methods

### Preparation of Syndecan-4 Proteoliposomes

To create liposomes, 10 mg/ml stock solutions of 1,2-dioleoyl-sn-glycero-3-phosphocholine, 1,2-dioleoyl-sn-glycero-3-phosphoethanolamine, sphingomyelin, and cholesterol were prepared in chloroform. Stock lipid solutions were mixed in a round bottom flask at a volumetric ratio of 2:1:1:1, respectively. The chloroform was then removed using a rotary evaporator attached to a vacuum pump to create a dried lipid film. Liposomes were resuspended in HEPES buffered salt solution. The liposome solution was put through three cycles of sonication for 30 seconds, vortexing for 1 minute, freezing, and thawing. The liposomes are then extruded through a 400 nm polycarbonate filter. Octyl-b-glucopyranoside (OG) is added to the lipid solution to make a 1% w/v solution. A syndecan-4 protein solution was then prepared at 71 μg/ml. OG was added to the protein solution to create a 1% w/v mixture. The next steps were performed at 4°C. The lipid and protein solutions were mixed at 1:1 and left on a rocking shaker for 30 minutes. OG concentration was diluted by adding 10% volumes of PBS four times, leaving 30-minute intervals between each dilution. Dialysis was performed on the solution overnight against PBS at 300 kDa. Excess lipids were removed from the solution using Biobeads.

### Preparation of Alginate Gels and Crosslinker

Sterile alginate powder (Sigma) was added to sterile saline to create a 2% w/v solution. syndecan-4 proteoliposomes (2 μg/ml), FGF-2 (20 μg/ml), and/or PDGF-BB (100 μg/ml) were added to the alginate solution. To prepare the crosslinker, calcium sulfate was added to sterile saline to create 0.2% w/v solution. Both the alginate solution and the crosslinker were taken up into a 1 ml syringe just before injection (100 μl of each solution).

### Induction of Diabetes in Rabbits

Studies involving animals were performed with the approval of the University of Texas at Austin and the UTHealth Science Center at Houston Institutional Animal Care and Use Committee (IACUC), the Animal Care and Use Review Office (ACURO) of The United States Army Medical Research and Materiel Command Office of Research Protections, and in accordance with NIH guidelines for animal care. New Zealand rabbits were transitioned from standard alfalfa chow to a 0.1% cholesterol diet over the course of five days. After two weeks on the 0.1% cholesterol diet, rabbits were induced with diabetes using an intravenous alloxan injection. Briefly, the rabbits were sedated. A bassline blood glucose measurement was attained. Eight milliliters of alloxan at 100 mg/kg was injected through an IV into the rabbit over an eight-minute period using a syringe pump. Blood glucose levels were monitored closely for 12 hours after the injection. A successful induction of diabetes was determined if the rabbit’s blood glucose level remained over 150 mg/dl prior to insulin administration.

### Hind limb Ischemia Surgery and Treatments

To induce acute ischemia in the hind limb of New Zealand rabbits, a longitudinal incision was made in the skin over the femoral artery. The femoral artery was exposed using blunt dissection. One percent lidocaine was applied to the area to reduce nerve irritation and promote vasodilation. Continued blunt dissection was used to expose the entire length of the femoral artery and branches including the inferior epigastric, deep femoral, lateral circumflex, and superficial epigastric arteries. The tissue was kept moistened with saline to avoid damage. The femoral artery was then carefully separated from vein and nerve. The femoral artery was then ligated with 4.0 silk sutures, cut, and excised. Two weeks after hind limb ischemia surgery, ten syringes containing 100 μl of alginate with treatment and 100 μl of calcium sulfate crosslinker were prepared for intramuscular injection (200 μl total volume of injection). Treatments included S4PLs+FGF-2+PDGF-BB in alginate, S4PLs+FGF-2 in alginate, FGF-2 in alginate, and an alginate-only control.

### Angiography Quantification

Angiograms were quantified using grid analysis techniques with ImageJ Software. Briefly, brightness and contrast were adjusted to better visualize vessels. A grid overlay of 100 pixels per square was used for foot quantification and 750 pixels per square was used for the thigh vasculature quantification. The multi-point tool was used to count intersections between the grid and vessels. To access the carotid artery, an incision was made just lateral to the trachea. Blunt dissection was used to expose the carotid artery and separate it from the jugular vein and vagus nerve. Ligatures were placed at the proximal and distal ends of the carotid artery. The distal end was tied off and a ligaloop was placed at the proximal end. A wire insertion tool was then inserted into the artery. Using the tool, a guidewire was fed into the artery to the aortic bifurcation in the descending aorta. The insertion tool was then removed and a 3F pigtail angiographic catheter was placed over the wire and advanced 2 cm proximal to the aortic bifurcation. Nitroglycerine and lidocaine were given to increase vasodilation. Contrast media was injected through the catheter using an automated angiographic injector. Angiography was performed before femoral ligation, after femoral ligation, and before sacrifice at week 10.

### Toxicology/Histological Analysis

To assess potential toxic effects of the treatments, mice were injected with of the following treatments into their left thigh muscle: no treatment, FGF-2 only, PDGFBB only, S4PLs only, FGF-2 + S4PLs, PDGFBB + S4PLs, and FGF-2 + PDGFBB + S4PLs. The concentration of the drug was 20 µg FGF-2, 2 µg S4PL and/or 100 µg PDGF-BB in 100 µL. This is roughly 100x the dose used in rabbits, normalized to body weight. The mice were monitored for 14 days after injection for signs of stress, weight loss, and/or pain. On day 14, the mice were sacrificed and underwent perfusion fixation to preserve the tissues. The liver, lung, spleen, pancreas, heart, kidney, thigh muscle, and aorta were all harvested for histological analysis.

### Stability testing for syndecan-4 proteoliposomes

Syndecan-4 proteoliposomes were stored under argon gas and inside a secondary vacuum sealed bag. The treatments were added to human umbilical vascular endothelial cells (HUVECs) for 30 minutes. The cells were then washed with PBS and lysed in the following lysis buffer: 20 mM Tris, 150 mM NaCl, 1% Triton X-100, 0.1% SDS, 2 mM sodium orthovanadate, 2 mM PMSF, 50 mM NaF, and protease inhibitors (Roche). The lysates were sonicated three times for 1 min and kept on ice in between sonication bouts. Total protein was measured using a BCA assay (Thermo Scientific) and equal amounts of protein were loaded per lane. SDS page was performed on the samples and the proteins were transferred to nitrocellulose membranes using wet transfer (Bio-Rad Laboratories). The membranes were blocked with PBS with 0.1% Tween-20 (PBST) with 5% Blotto for one hour and then incubated with antibodies to phospho-p90RSK (Thr359/Ser363; #9344; Cell Signaling Technology, Inc.) and RSK1/RSK2/RSK3 (32D7; #9355; Cell Signaling Technology, Inc.) overnight at 4°C. The membranes were then incubated with HRP conjugated secondary antibodies (Santa Cruz) at 1:3500 dilution in PBST with 1% Blotto for two hours at room temperature. The membranes were then exposed to ECL solution (Super Signal West Femto; Thermo Scientific) and imaged with a digital imaging system (G:Box Chemi XX6; Syngene).

### Numerical Simulation

Comsol Multiphysics (version 5.6; Comsol, Inc., Burlington, MA) was used to simulate the diffusion of FGF-2 through the tissue. The FGF-2 was assumed to have a diffusivity of 6 × 10^−9^ cm^2^/s in the tissue. Based on the actual injection volume and concentration, there was a total of 12.2 µg of FGF-2 per injection that was assumed to be completely released over 7 days. The geometry of the tissue was assumed to be as shown in **Supplemental Fig. 1**, to mimic the injections in the rabbit hindlimb muscles. The injections were assumed to create a sphere of the volume of the injection.

**Figure 1.**
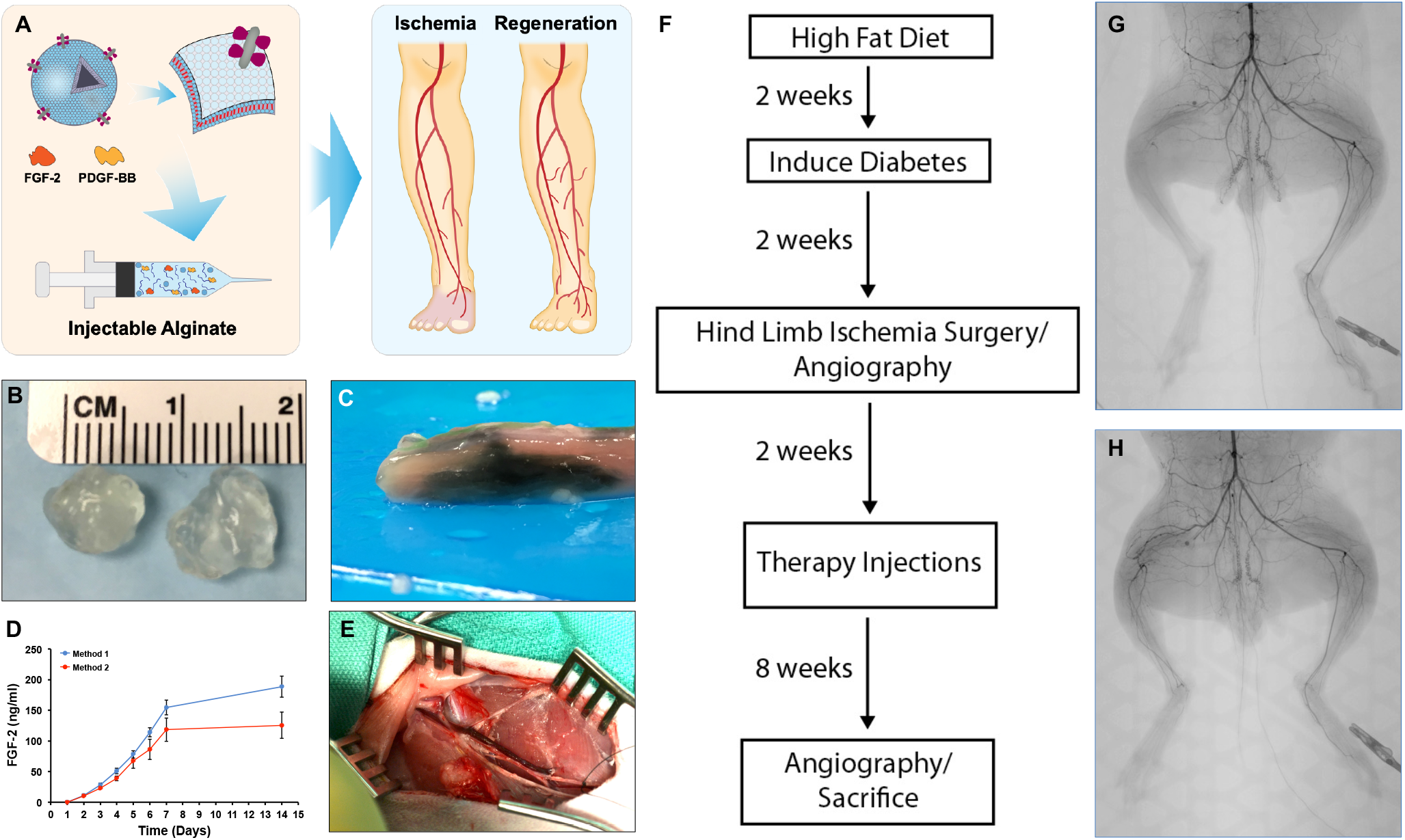
Treatment and Experimental Design. (A) Diagram of the therapeutic agents and their use in treating peripheral ischemia. Syndecan-4 proteoliposomes are combined with FGF-2 and PDGF-BB and then mixed with an injectable alginate formulation. These compounds are injected into ischemic regions to induce revascularization of the tissue. (B) The controlled release alginate after gelation. (C) The alginate was combined with a blue dye and injected into muscle tissue. The muscle was then cut to visualize the intramuscular distribution of the alginate after injection. (D) Release profile of FGF-2 from the alginate gel using two different formulations. Method 2 was chosen for further studies. (E) The surgical field during the femoral ligation surgery in diabetic, hyperlipidemic rabbits. (F) Overall procedure for the establishment of hyperlipidemia/diabetes, surgical ligation and analysis of the rabbits. (G, H) Representative angiograms before (G) and after(H) surgical ligation of the femoral artery.

### Statistical Analysis

All results are shown as mean ± standard error of the mean. Comparisons between only two groups were performed using a 2-tailed Student’s t-test. Differences were considered significant at p<0.05. Multiple comparisons between groups were analyzed by 2-way ANOVA followed by a Tukey post-hoc test. A 2-tailed probability value <0.05 was considered statistically significant.

## Results

### Development of a gel formulation of syndecan-4 proteoliposomes for intramuscular injection

We created and optimized and alginate gel for intramuscular injection (**Fig. 1A, B**). Using a dye impregnated gel we found that the gel distributed in the muscle on injection (**Fig. 1C**). The compounds were almost completely released after seven days (**Fig. 1D**). Rabbits were given a high fat diet for 4 weeks and made diabetic for 2 weeks prior to surgery (**Fig. 1E**). To create limb ischemia, the femoral artery was ligated in the right leg and the rabbits were allowed to recover for two weeks to avoid treatment during the acute healing phase of recovery (**Fig. 1F**). Our group has previously optimized this model and shown that it has significantly impaired revascularization compared to control rabbits and other models.^14^ The rabbits were given ten injections of alginate with the treatments and recovery was tracked using angiography (**Fig. 1G, H**).

### Syndecan-4 proteoliposomes with growth factors enhance revascularization of the ischemic thigh and calf muscles of diabetic, hyperlipidemic rabbits

Angiograms were taken before, immediately after, and ten weeks after hind limb ischemia surgery. Immediately following the procedure, no vessels were visible past the mid-thigh in the ischemic limb. After ten weeks, vascular recovery was assessed across multiple treatment groups using grid intersection analysis. Briefly, a grid was superimposed onto the angiograms and intersections between the grid and vessels were counted. These numbers were either reported as an intersection count or ratioed to create a relative vascularity metric. Overall, treatments without S4PLs showed some vessels extending from the proximal thigh to the distal thigh compared to treatments including S4PLs which appeared to have more robust vascular network formation with more densely packed new vessel growth. Grid intersection counts showed significant increases in vasculature for the Alg+S4PLs+FGF-2 treatment group compared to the alginate control and FGF-2 groups (**Fig 2A, B**; **Supplemental Fig. 2**). When ratioed to the contralateral control limb at week 10, the Alg+PDGFBB+S4PLs+FGF-2 group significantly improved recovery over all other groups including Alg+S4PLs+FGF-2 (**Fig. 2C**). A ratio was also taken of the grid intersections counted in the ischemic thigh at week 10 to the intersections counted in the ischemic thigh before hind limb surgery. These results show a significant increase in vascularity for the Alg+PDGFBB+S4PLs+FGF-2 group as well (**Fig 2D**). A count was taken of new vessel growth, indicated by tortuosity, which showed a significant increase for both S4PL-containing treatments compared to the alginate control and FGF-2 groups. We performed also performed the model on one rabbit using Alg+PDGFBB+FGF-2 and found a similar response to the FGF-2 only and alginate groups (**Supplemental Fig. 3**).

**Figure 2.**
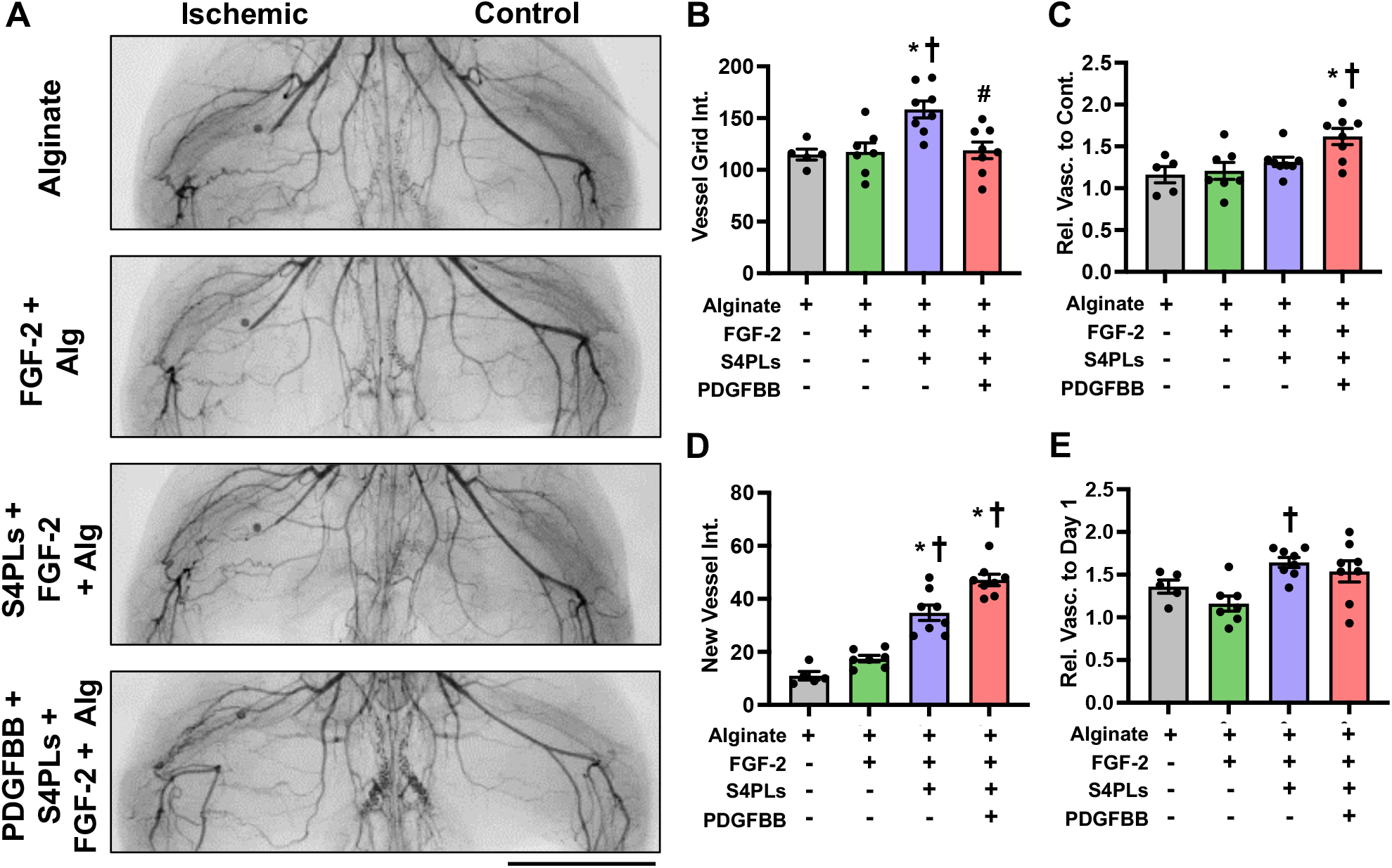
Syndecan-4 proteoliposomes as a co-treatment enhances growth factor induced revascularization in the thigh of diabetic, hyperlipidemic rabbits with hindlimb ischemia. (A) Angiograms of the thighs of the rabbits prior to sacrifice. The control limb is shown on the right and the limb with the ligated femoral artery is on the left. Scale bar = 5 cm. (B) Quantification of vessels counted in the ischemic thigh at the model endpoint. The vessels are counted as number of intersections with an overlayed grid. (C) Relative vascularity of the ischemic thigh ratioed to the contralateral control thigh at the model endpoint. (D) Quantification of new vessels counted in the ischemic thigh at the model endpoint. (E) Relative vascularity of the ischemic thigh ratioed to the thigh at day 1 prior to ligation. \**p* < 0.05 vs Alginate. ^†^*p* < 0.05 vs. FGF-2 + Alg. ^#^*p* < 0.05 vs S4PLs + FGF-2 + Alg. The group sizes were as follows: Alginate (n = 5), FGF-2 + Alg (n = 7), S4PLs + FGF-2 + Alg (n = 8), and S4PLs + FGF-2 + PDGF-BB + Alg (n = 8).

**Figure 3.**
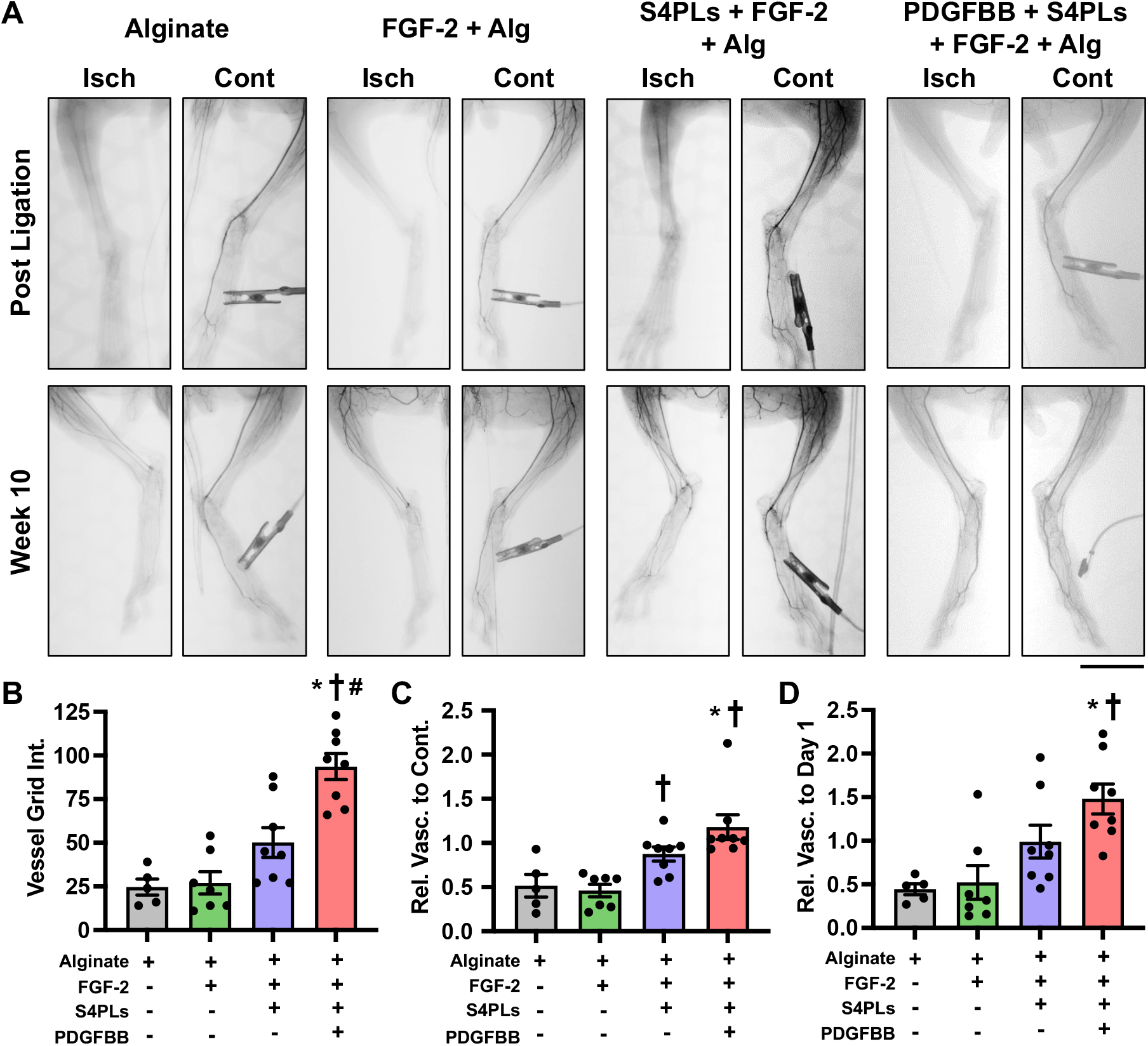
Syndecan-4 proteoliposomes enhance growth factor induced revascularization in the calf and foot of diabetic, hyperlipidemic rabbits with hindlimb ischemia. (A) Angiograms of the lower limb of the rabbits prior to sacrifice. Isch. = ischemic limb; Cont. = control limb. Scale bar = 5 cm. (B) Quantification of vessels counted in the ischemic calf at the model endpoint. The vessels are counted as number of intersections with an overlayed grid. (C) Relative vascularity of the ischemic limb ratioed to the contralateral control limb at the model endpoint. (D) Relative vascularity of the ischemic thigh ratioed to the thigh at day 1 prior to ligation. \**p* < 0.05 vs Alginate. †*p* < 0.05 vs. FGF-2 + Alg. ^#^*p* < 0.05 vs S4PLs + FGF-2 + Alg. The group sizes were as follows: Alginate (n = 5), FGF-2 + Alg (n = 7), S4PLs + FGF-2 + Alg (n = 8), and S4PLs + FGF-2 + PDGF-BB + Alg (n = 8).

### Syndecan-4 proteoliposomes with growth factors enhance revascularization of the ischemic foot of diabetic, hyperlipidemic rabbits

Angiograms were taken before, immediately after, and ten weeks after hind limb ischemia surgery. Immediately following the procedure, no vessels were visible in the ischemic foot (**Fig. 3A**). After ten weeks, differing levels of vascular recovery were observed across treatment groups. Treatments without S4PLs showed small vessels extending from the ankle into the foot. Midway into the foot toward the toes, little to no vascular recovery was observed. Treatments including S4PLs had a more robust vascular network formation with vessels clearly extending into the toes of the rabbits. Furthermore, the Alg+PDGFBB+S4PLs+FGF-2 group showed larger vasculature that extended farther down into the foot. A grid was superimposed onto the angiograms and intersections between the grid and vessels were counted. The Alg+PDGFBB+S4PLs+FGF-2 treatment group produced significantly more grid intersections compared to the alginate control and FGF-2 groups (**Fig. 3B**). When ratioed to the contralateral control limb at week 10, groups containing S4PLs significantly improved recovery over the alginate control and FGF-2 groups (**Fig. 3C**). A ratio was also taken of the grid intersections counted in the ischemic foot at week 10 to the intersections counted in the ischemic foot before hind limb surgery. These results show a significant increase in vascularity for the Alg+PDGFBB+S4PLs+FGF-2 group as well (**Fig. 3D**). Blood pressure measurements were taken using a cuff placed at the ankle. Here, all treatments showed an improved ratio at week 10 over the alginate-only control (**Fig. 3E**).

### Syndecan-4 proteoliposomes with growth factors increased vascularity of the ischemic hindlimb of rabbits with diabetes and hyperlipidemia

We next performed a histological analysis on biopsies from the ischemic muscle of the rabbits (**Fig. 4A**). Immunostaining for PECAM on the sections demonstrated increased small blood vessels in the Alg+S4PL+FGF-2 and Alg+PDGFBB+S4PLs+FGF-2 groups (**Fig. 4B**). In addition, found an increase in arterioles in the Alg+S4PL+FGF-2 and Alg+PDGFBB+S4PLs+FGF-2 groups (**Fig. 4C**).

**Figure 4.**
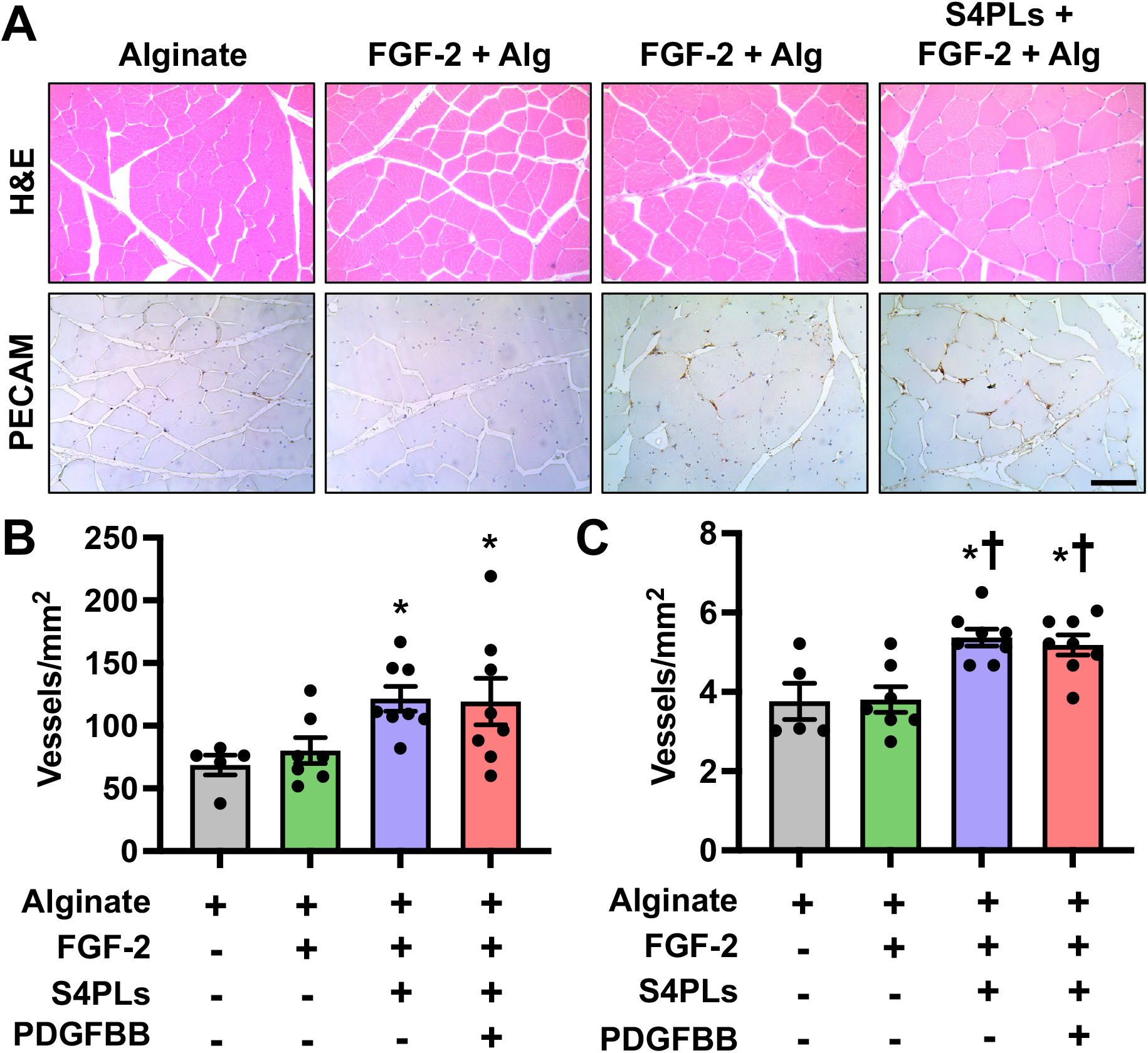
Syndecan-4 proteoliposomes enhance vascularity in the ischemic limbs of hyperlipidemic rabbits. (A) Histological analysis of biopsies from the ischemic hind limbs of rabbits. Scale bar = 100 µm. (B) Quantification of small vessels counted in the ischemic calf at week 10. (C) Quantification of arterioles vessels counted in the ischemic calf at week 10. *p < 0.05 versus alginate group. ^†^p < 0.05 versus the FGF-2 group. The group sizes were as follows: Alginate (n = 5), FGF-2 + Alg (n = 7), S4PLs + FGF-2 + Alg (n = 8), and S4PLs + FGF-2 + PDGF-BB + Alg (n = 8).

### Stability and Toxicity of Syndecan-4 Proteoliposomes

Stability in storage and transport is essential for therapies to be used in a hospital environment. The liposome carrier in S4PLs is not stable to freezing and thus requires storage under refrigeration or at room temperature. We tested the stability of S4PL by storing them under argon gas at 4°C or 20°C for varying time periods out to 28 days. The S4PLs were combined with FGF-2 and tested for activity by stimulating endothelial cells and assess the activation of the RSK pathway. S4PLs stored at 4°C were stable and maintained their ability to enhance FGF-2 activity for at least 28 days (**Fig. 5A, B**). To assess potential toxic effects of syndecan-4 proteoliposomes, we injected high concentrations of the treatments into the left thigh of mice. The following treatments were injected: FGF-2, PDGF-BB, S4PLs, FGF-2 + S4PLs, PDGFBB + S4PLs, and FGF-2 + PDGFBB + S4PLs. All of the treatments were injected at 100x the concentration used in rabbits (adjusted for body weight). The mice were monitored for 14 days after injection but no signs of stress, weight loss, or pain. On day 14, the mice were sacrificed and the tissues harvested for histological analysis. There were no signs of toxicity in any of the organs from the groups (**Fig. 5C**).

**Figure 5.**
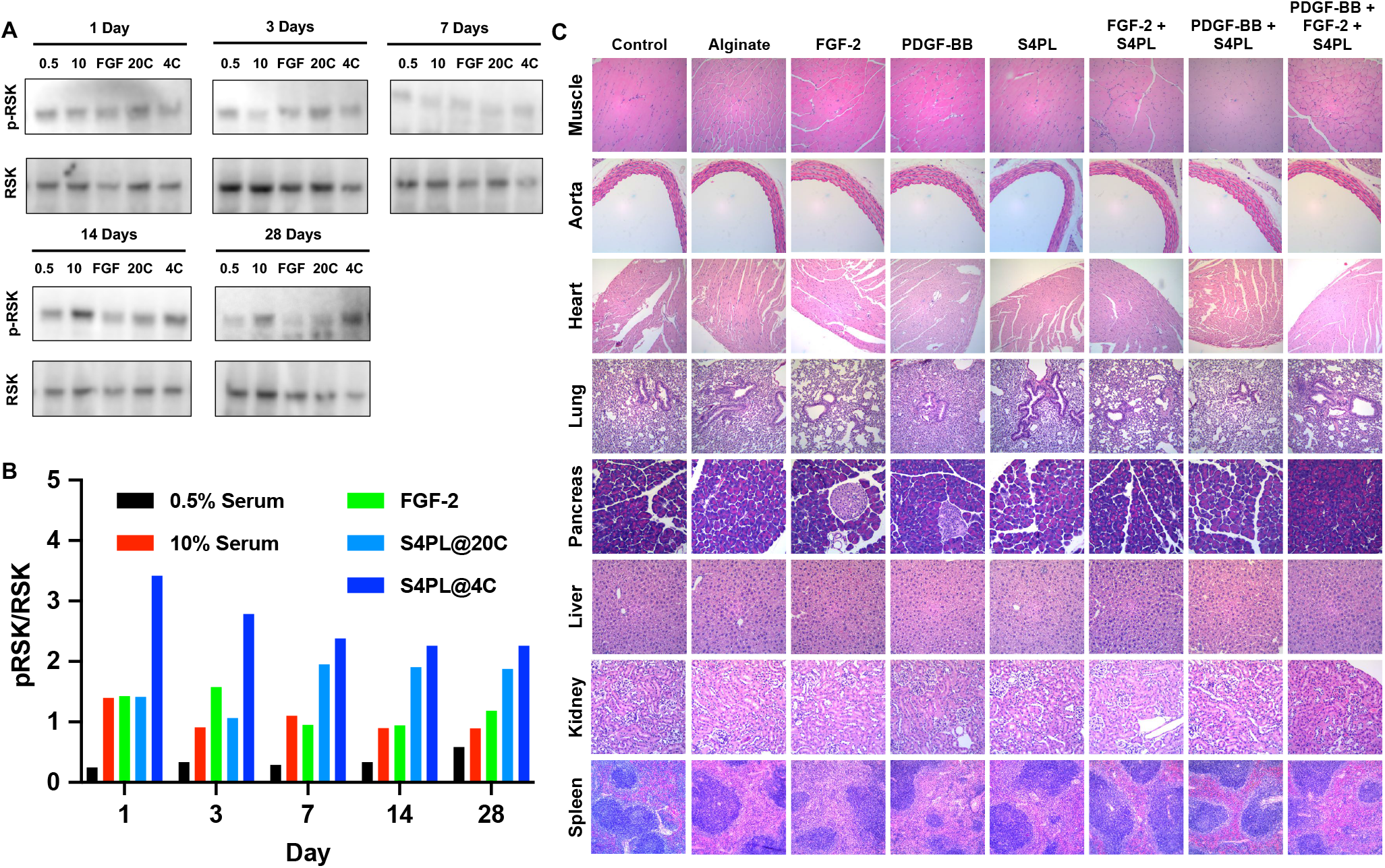
Ex vivo stability and toxicity testing in mice for syndecan-4 proteoliposome treat. (A) Syndecan-4 proteoliposomes were stored at either 20°C or 4°C for the times indicated. At the time of testing, the syndecan-4 proteoliposomes were combined with 10 ng/ml FGF-2. Endothelial cells were treated with the indicated treatments for 30 minutes, lysed and then immunoblotted for pRSK or RSK. The treatments were as follows: 0.5 = 0.5% FBS, 10 FBS, FGF-2 = 10 ng/ml FGF-2, S4PL@20C = syndecan-4 proteoliposomes stored at 20°C + 10 ng/ml FGF-2, and S4PL@4C = syndecan-4 proteoliposomes stored at 4°C + 10 ng/ml FGF-2. (B) Ratio of densimetry of western blots for pRSK and RSK. (C) Histology analysis of tissue from mice treated with 20 µg FGF-2, 2 µg S4PL and/or 100 µg PDGF-BB in 100 µL.

### Computational model of growth factor delivery from intramuscular injections

To understand the release kinetics and distribution of the treatments, we created a computational model of the release within the hindlimb of the rabbit. In the model, we assumed that the injection forms sphere within the tissue and releases drug at the rate that is the same as the ex vivo release studies. We created a model of the thigh muscle of the rabbit with the injections in an array similar to our injections in the animal model. As uptake and degradation are difficult to estimate accurately within the tissue and not included, the model represents the total delivery of drug did the tissue rather than the actual concentration. The release simulation showed increasing drive overtime within the tissue that reached approximately 25 µg/cm^3^ of drug delivered in the region between the spheres on each line of injections at the lowest point over 7 days (**Fig. 6A, B**). In the region between the lines of injections, the minimum drug delivered was approximately 10 µg/cm^3^ over 7 days.

**Figure 6.**
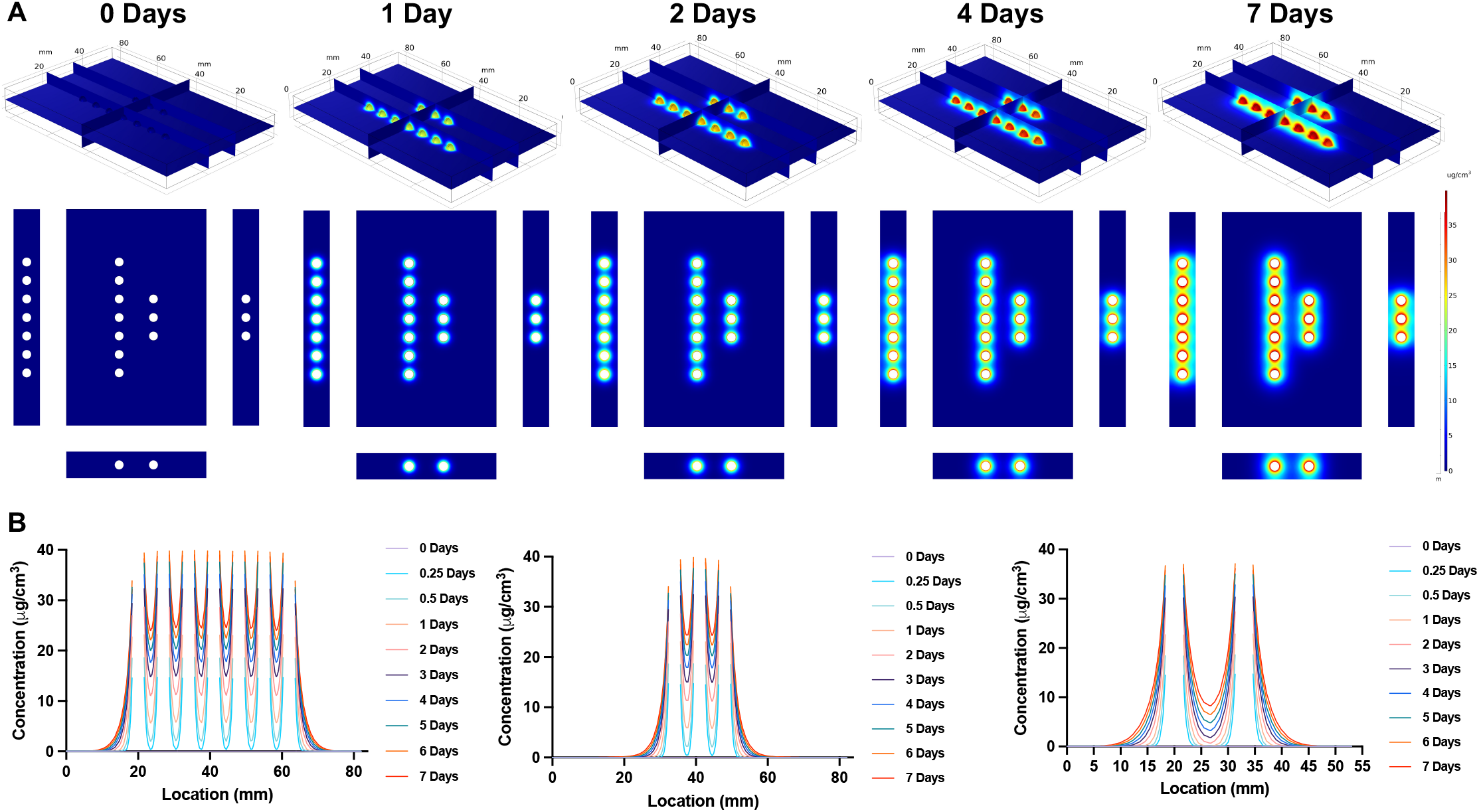
Computational model of growth factor delivery from alginate gels injected into the rabbit limb. (A) Computational model of the release of FGF-2 from alginate gels in an array matching the injections in the rabbit leg. The model approximates the size and location of the intramuscular injections. (B) Line scans of concentration averaged over the muscle along the vertical lines shown in part A.

## Discussion

The enhancement of revascularization of ischemic tissues using regenerative therapies has been a major goal in cardiovascular medicine. While the concept of therapeutic angiogenesis/arteriogenesis is highly appealing, multiple therapies in this area have failed to produce significant benefits for patients in clinical trials.^6,20,21^ In contrast, many preclinical studies in large animals demonstrate benefits from regenerative therapies in terms enhancing vascularization.^22-24^ In this work, we tested the efficacy of growth factors delivered from an alginate gel in combination with S4PLs in a rabbit model of hindlimb ischemia with diabetes and hyperlipidemia. Our group has previously optimized this animal model to produce therapeutic resistance to growth factor therapies and lasting ischemia beyond similar models in healthy rabbits.^14^ Our previous work has demonstrated that S4PLs in combination with growth factors can enhance revascularization in healthy and diabetic mice, healthy rats and wound healing in diabetic mice.^9-12,19^ In addition, we have shown that mechanistically S4PLs enhance FGF-2 activity in endothelial cells, alter subcellular trafficking of FGF-2 including increasing uptake and recycling of the growth factor,^12^ and can increase the numbers of M2 macrophages in healing wounds in diabetic mice.^10,12^ This study furthers the translational development of this therapeutic by demonstrating that S4PLs in combination with growth factor delivery produces marked enhancement of revascularization in the ischemic hindlimb of rabbits in this model. In addition, we have shown that S4PLs are stable under refrigeration conditions for at least 28 days, enabling the logistics for delivery of the compounds to health care providers for clinical use. Finally, we have developed a computational model of the delivery of the treatments during local delivery to help to guide clinical application.

The translational disconnect between preclinical and clinical results suggests an inadequacy in predictive value of limb ischemia models performed in healthy rabbits for determining outcomes in treating ischemia in patients with PAD. The animal model that was used in our studies was previous optimized to have prolonged recovery from ischemia and resistance to growth factor induction of revascularization.^14^ In this study, FGF-2 alone did not provide enhanced revascularization as assessed by angiography or histological analysis. This finding is in contrast to multiple studies that have found enhancement of revascularization in the ischemic hindlimbs of healthy rabbits using FGF-2.^25-32^ The results from the advanced rabbit model are consistent with the lack of efficacy for FGF-2 treatments in clinical trials for intermittent claudication^8^ and suggests that the optimized rabbit model may provide better correlation with clinical outcomes.

In our study, the addition of S4PLs markedly improved the efficacy of growth factors in the rabbit ischemia model. On histology, both the S4PL+FGF-2 and S4PL+FGF-2+PDGFBB groups had improvements in both small capillaries and larger arterioles. However, on angiographic analysis the PDGF-BB containing group had the greatest increases in vascularity. This suggests benefits for including PDGF-BB in the therapy, particularly for the larger vessels that arise from arteriogenesis. Previous studies have supported a similar result, demonstrating that PDGF-BB can act synergistically with FGF-2 to enhance angiogenesis and facilitate stabilization of blood vessels.^24,28^ Notably, our previous also supports that S4PLs can enhance PDGF-BB activity toward diabetic wound healing in the absence of exogenously delivered FGF-2.^9^ While it was not feasible for us to test all combinations of the therapy components the animal model, we did perform the model in one rabbit with FGF-2 and PDGF-BB. This rabbit showed similar revascularization to the alginate and alginate with FGF-2 groups. Due to a lack of standardization in the methods used in the performance and analysis of preclinical models of hindlimb ischemia in rabbits, direct comparison to other treatments is challenging. In general, the enhancement in revascularization seen in our study in this disease rabbit model is equivalent or greater than previous studies using healthy rabbit models of hindlimb ischemia to evaluate protein therapeutics,^7,8,22-42^ gene therapy,^23,43-55^ or cell therapies.^56-60^ Thus, our work supports that S4PL provides marked benefits for enhanced revascularization in comparison to growth factors alone and shows promise in addressing therapeutic resistance in pro-angiogenic treatments.

In conclusion, our work has shown that S4PLs combined with angiogenic growth factor have efficacy in a large animal model of limb ischemia with diabetes and hyperlipidemia. These treatments work despite the therapeutic resistance present in this model and thus may have promise as effective therapies for ischemia in peripheral arterial disease. Local delivery with an optimized system allows sustained, targeted treatment for ischemic regions, likely a more effective strategy than systemic or non-sustained release methods for growth factor delivery. In addition, our studies suggest that this treatment has low toxicity even elevated concentrations. Overall, the treatments have promise for enabling vascular regeneration in patients with ischemia.

## Acknowledgements

The authors gratefully acknowledge funding through the DOD CDMRP (W81XWH-16-1-0580; W81XWH-16-1-0582) and the National Institutes of Health (1R01HL141761-01) to ABB.

## Supplemental Figures

**Supplemental Figure 1.**
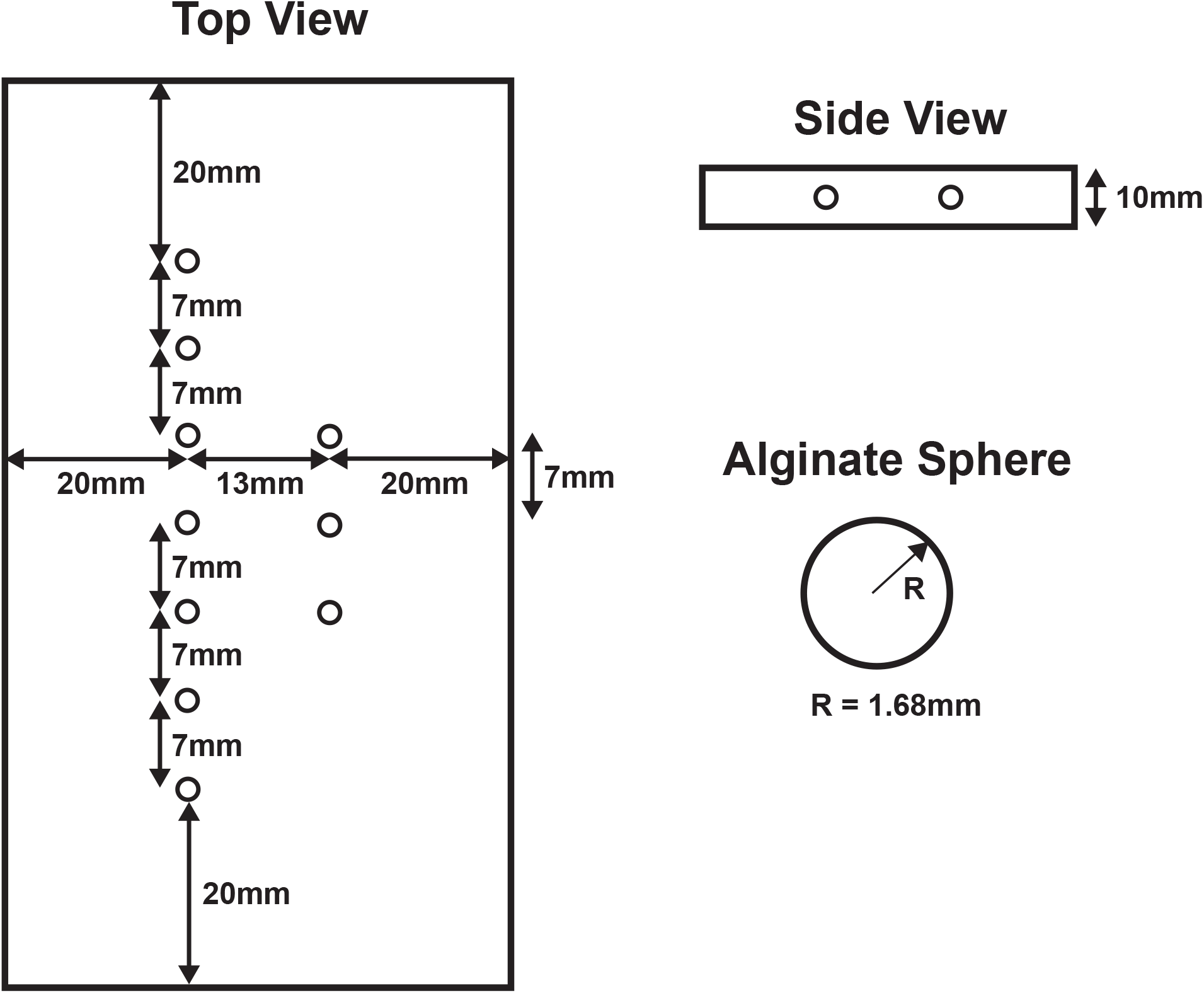
Geometry used for the simulation of drug diffusion in the tissue.

**Supplemental Figure 2.**
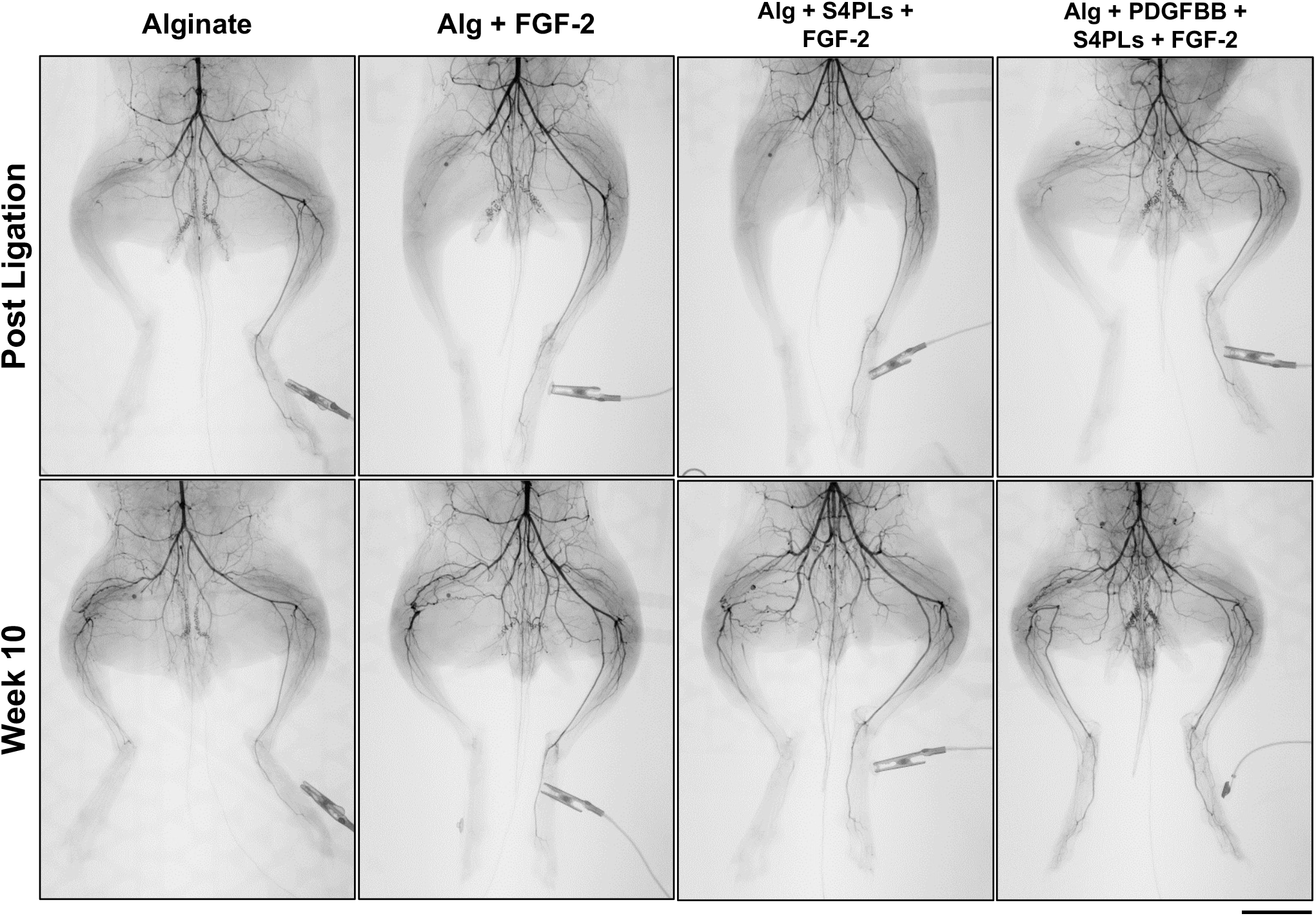
Full angiograms for the rabbits post-ligation and after 10 weeks. Bar = 5 cm.

**Supplemental Figure 3.**
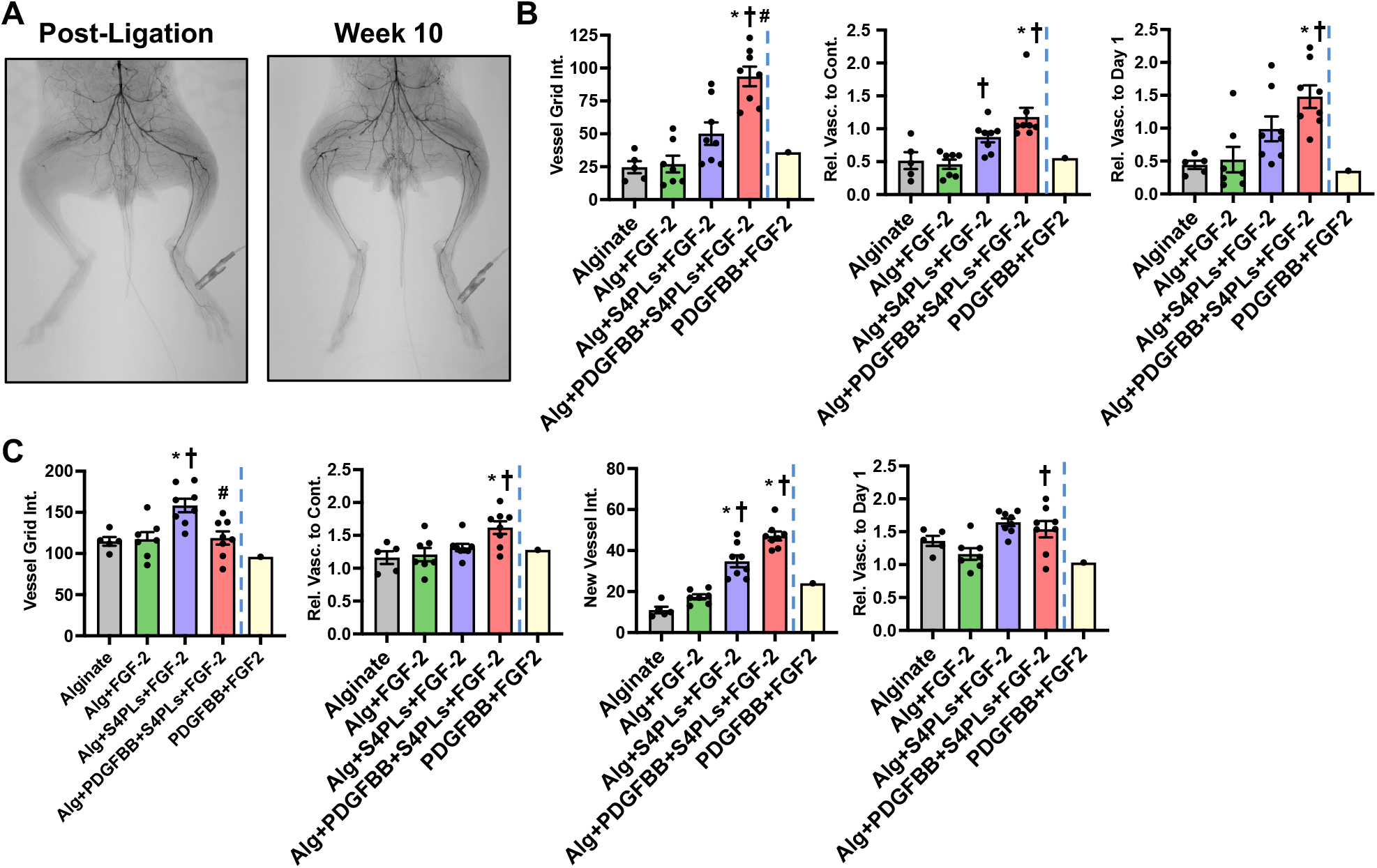
(A) Angiography from rabbit treated with PDGF-BB+FGF2. (B) Summary of data of the study in comparison a rabbit treated with PDGF-BB+FGF2 (shown in yellow).

